# Molecular Weight-dependent Diffusion, Biodistribution, and Clearance Behavior of Tetra-armed Poly(ethylene glycol) Subcutaneously Injected into the Back of Mice

**DOI:** 10.1101/2023.03.08.531818

**Authors:** Shohei Ishikawa, Motoi Kato, Jinyan Si, Lin Chen Yu, Kohei Kimura, Takuya Katashima, Mitsuru Naito, Masakazu Kurita, Takamasa Sakai

## Abstract

Four-armed poly(ethylene glycol) (PEG)s are essential hydrophilic polymers extensively utilized to prepare PEG hydrogels, which are valuable tissue scaffolds. When hydrogels are used *in vivo*, they eventually dissociate due to the cleavage of the backbone structure. When the cleavage occurs at the cross-linking point, the hydrogel elutes as an original polymer unit, i.e., four-armed PEG. Although four-armed PEGs have been utilized as subcutaneously implanted biomaterials, the diffusion, biodistribution, and clearance behavior of four-armed PEG from the skin are essential. This paper investigates time-wise diffusion from the skin, biodistribution to distant organs, and clearance of fluorescence-labeled four-armed PEGs with molecular weight (*M_w_*) ranging from 5–40 kg/mol subcutaneously injected into the back of mice. Changes over time indicated that the fate of subcutaneously injected PEGs is *M*_w_-dependent. Four-armed PEGs with *M*_w_ ≤ 10 kg/mol gradually diffused to deep adipose tissue beneath the injection site and distributed dominantly to distant organs, such as the kidney. PEGs with *M*_w_ ≥ 20 kg/mol stagnated in the skin and deep adipose tissue, and were mainly delivered to the heart, lung, and liver. The fundamental understanding of the *M*_w_-dependent behavior of four-armed PEGs is beneficial for preparing biomaterials using PEGs, providing a reference in the field of tissue engineering.

## Main text

Poly(ethylene glycol) (PEG)s are representative hydrophilic and biocompatible polymers utilized to synthesize biomaterials such as artificial extracellular matrix (ECM), drug delivery carriers, and hydrogels because of their bioinert nature originating from high volume exclusion, low immune reaction, and protein exclusion.^1–4^ Owing to their unique characteristics, COVID- 19 vaccines include PEGs with a molecular weight (*M_w_*) of approximately 2 kg/mol, which is widely shared and studied by several scientists and clinicians, indicating an increasing focus on PEG application and characteristics.^5–7^ Drugs comprising or conjugating with PEGs, such as PEGylated materials, administered intravenously or intramuscularly *in vivo*, are first distributed to organs such as the heart, lungs, liver, and kidney before being excreted from the body through the clearance system.^8–10^ The biodistribution and excretion of molecules intravenously injected can be affected by polymer *M*_w_, molecular structure, charge, and hydrophilic/hydrophobic properties.^8,11–14^ Therefore, establishing the basis of biodistribution of polymers *in vivo* has facilitated the development of interferon, vaccine dispersant, and PEGylated drugs.^15^ Intelligent polymer micelles consisting of PEG copolymers have been developed to accumulate in tumors via enhanced permeability and retention effects. Moreover, methods for establishing controlled drug delivery systems have been proposed, resulting in clinical trials.^16^ This success is due to the extensive research on the biodistribution and clearance of polymers by researchers worldwide. Therefore, basic information regarding biodistribution and excretion is critical for the development of polymeric biomaterials for clinical implementation.

Although several reports have extensively examined the fate of intravenously-injected polymeric composites, the clearance behavior of PEGs injected subcutaneously remains unclear. Four-armed PEG, in particular, functions as a fundamental polymer to produce biomaterials and is thus suitable for preparing PEG-based hydrogels,^17,18^ which serve as hemostatic agents, artificial vitreous, and cell scaffolds.^19–22^ When PEG hydrogels are implanted subcutaneously and the bond between four-armed PEGs is cleaved by chemical or physical stimuli, the original four-armed PEG is obtained as an eluate. A successful design control degradability; when tetra-PEG gels prepared using four-armed PEG with a *M*_w_ of 10 kg/mol is dissociated, four-armed PEG with a *M*_w_ of 10 kg/mol is ideally obtained. However, the biodistribution and clearance behavior of subcutaneously-injected four-armed PEG, especially its *M*_w_-dependence, is still elusive, limiting its clinical application. This information provides a concrete reference for using PEG hydrogels as biomaterials.

In this study, we characterized the diffusion, biodistribution to distant organs, and clearance behavior of four-armed PEG with a *M*_w_ of 5–40 kg/mol subcutaneously injected into the back of mice. Red fluorescence was used to partly functionalize four-armed PEG and visualize PEG localization, which was evaluated by fluorescence imaging analyses. *In vitro* permeation studies, in which a cell culture insert was set to a well plate, revealed the *M*_w_-dependence permeation behavior. Time-wise changes in the diffusion behavior from the injection site and in dissected deep adipose tissue beneath the injection site were characterized by observing fluorescence intensity at 0, 1, 4, 24, 48, 72, and 168 h after subcutaneous injection. Furthermore, biodistribution to distant organs, such as the heart, lung, liver, and kidney, was evaluated by dissecting these organs and observing fluorescence intensity, which suggested a *M*_w_-dependent biodistribution. Our findings indicate that PEG with a *M*_w_ below 10 kg/mol would be suitable precursors of hydrogels implanted subcutaneously because it readily diffuses once dissociated in subcutaneous tissue.

In this study, amine-terminated four-armed PEGs with a *M*_w_ of 5, 10, 20, and 40 kg/mol (hereafter referred to as PEG5, PEG10, PEG20, and PEG40, respectively) were partly modified using Alexa Fluor 594 (Alexa) through a condensation reaction between primary amine of PEG and *N*-hydroxysuccinimide ester (NHS) of Alexa to track diffusion, biodistribution, and clearance behavior of the four-armed PEGs (**Figure 1a**). The modification rate of Alexa to the end of the PEG was theoretically set to 0.1% vs. primary amine. The relative fluorescence intensity of a 10 g/L PEG solution in neutral phosphate-buffered saline (PBS) was similar to that of Alexa, even though the *M*_w_ differed (**Table S1**), indicating that the fluorescence was successfully tethered to PEGs regardless of the *M*_w_. Furthermore, since the amide cross-linking between PEG and Alexa hardly dissociates under *in vivo* conditions (pH 7.4 and 37 °C),^23^ fluorescence showed PEG localization, not that of Alexa molecules cleaved from PEGs.

**Figure 1.**
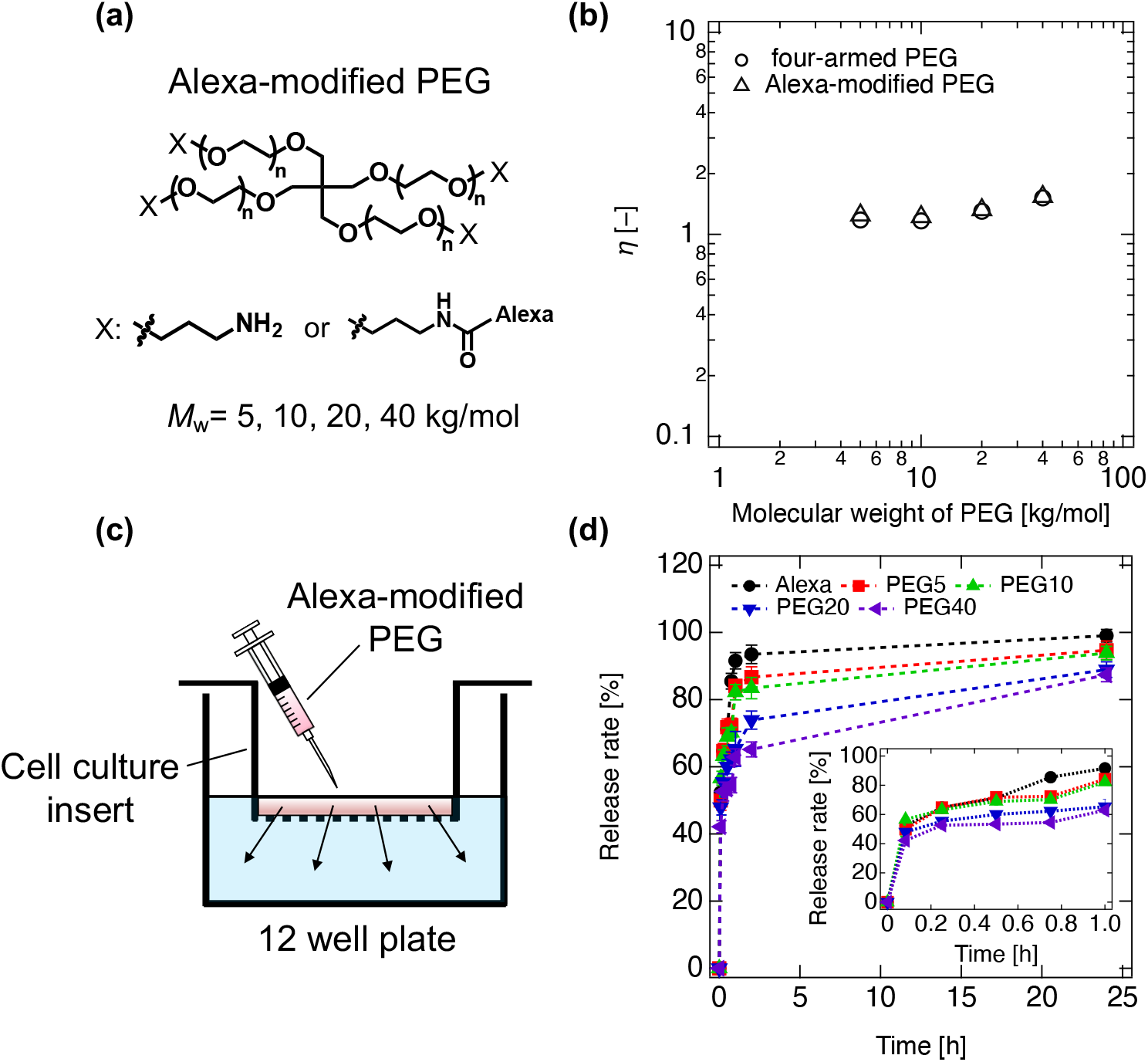
Characterization of Alexa-modified PEGs. **(a)** Conceptual chemical structure of four-armed PEGs. Four-armed PEGs functionalized with propyl amine were modified using Alexa Fluor 594 (referred to as Alexa). The molecular weight (*M*_w_) of PEGs was 5, 10, 20, and 40 kg/mol, thereafter denoted as PEG5, PEG10, PEG20, and PEG40, respectively. **(b)** Viscosity (*η*) relative to PBS as a function of *M*_w_. Circle and triangle symbols represent the *η* of PEGs and Alexa-modified PEGs, respectively. **(c)** Diagram of release behavior after adding the Alexa-modified PEG solutions. **(d)** Release rate of Alexa (black circle) and Alexa-modified PEG5 (red square), PEG10 (green triangle), PEG20 (blue inverted triangle), and PEG40 (purple left-arrowed triangle) from the top to the bottom compartment. Insert image shows release rate up to 1 h.

It is well-known that PEG can form nanoparticles when modified with hydrophobic fluorescence molecules.^12^ We confirmed that Alexa-modified PEGs did not form nanoparticles under experimental conditions based on viscosity and permeation measurements (**Figure 1b-d**). The viscosity of Alexa-modified PEG solutions was nearly the same as those of original PEG solutions regardless of the *M_w_* (**Figure 1b**), hinting that the modification did not induce polymeric nanoparticles. The slight increase in relative viscosity with *M*_w_ is likely due to an increase in hydrodynamic radius.^24,25^ Moreover, we investigated the permeation behavior of PEG using a membrane with a pore size of 3.0 μm (**Figure 1c** and **d**). The permeation rate decreased with increasing *M*_w_, and *M*_w_-dependent permeation was observed for 24 h. After 24 h, the release rate was 99.0%, 94.7%, 93.8%, 88.9%, and 87.4% for Alexa, PEG5, PEG10, PEG20, and PEG40, respectively. This *M*_w_-dependent permeation strongly suggests that the PEGs were successfully modified by Alexa, and PEG nanoparticles was negligible. Therefore, Alexa-modified four-armed PEG most probably exists as a polymer, allowing a detailed discussion on the *M*_w_-dependence, as detailed in the following sections.

To observe the clearance of PEGs from the subcutaneous tissue, we first subcutaneously injected a PBS solution of Alexa-modified PEGs in the back of mice (**Figure 2a**), and changes in local fluorescence intensity were evaluated by bidirectional imaging at 0, 1, 4, 24, 72, and 168 h post-injection (**Figure 2b**). From the top view, circular red fluorescence was observed on the initial injection site. With the decrease of fluorescence at the initial injection site, the fluorescence spread to nearby tissue, demonstrating diffusion of PEG molecules over time (**Figure 2b**). Quantitative analyses of local fluorescence images illustrated decreased injected PEG over time (**Figure 2c**) and the differences between PEGs of different *M*_w_. While PEG5 almost disappeared at 72 h, those with *M*_w_ > 5 kg/mol gradually attenuated over 168 h but did not completely disappear. The *M*_w_-dependent diffusion behavior *in vivo* was comparable to that of *in vitro* permeation, despite its diffusion speed being different. The diffusion behavior from the side view was consistent with the findings from the top view, showing a *M*_w_-dependent diffusion to deeper tissues (**Figure 2d-f**). Therefore, subcutaneously injected PEG molecules may diffuse toward deep tissues, such as deep adipose tissue beneath the injection site. Fluid in the skin and subcutaneous tissues was absorbed from collective lymph vessels or diffused to deep tissue, such as adipose tissues, transported into blood vessels via lymph nodes,^26,27^ and distributed to distant organs. It is reasonable to consider that macromolecules such as PEGs follow the same route.

**Figure 2.**
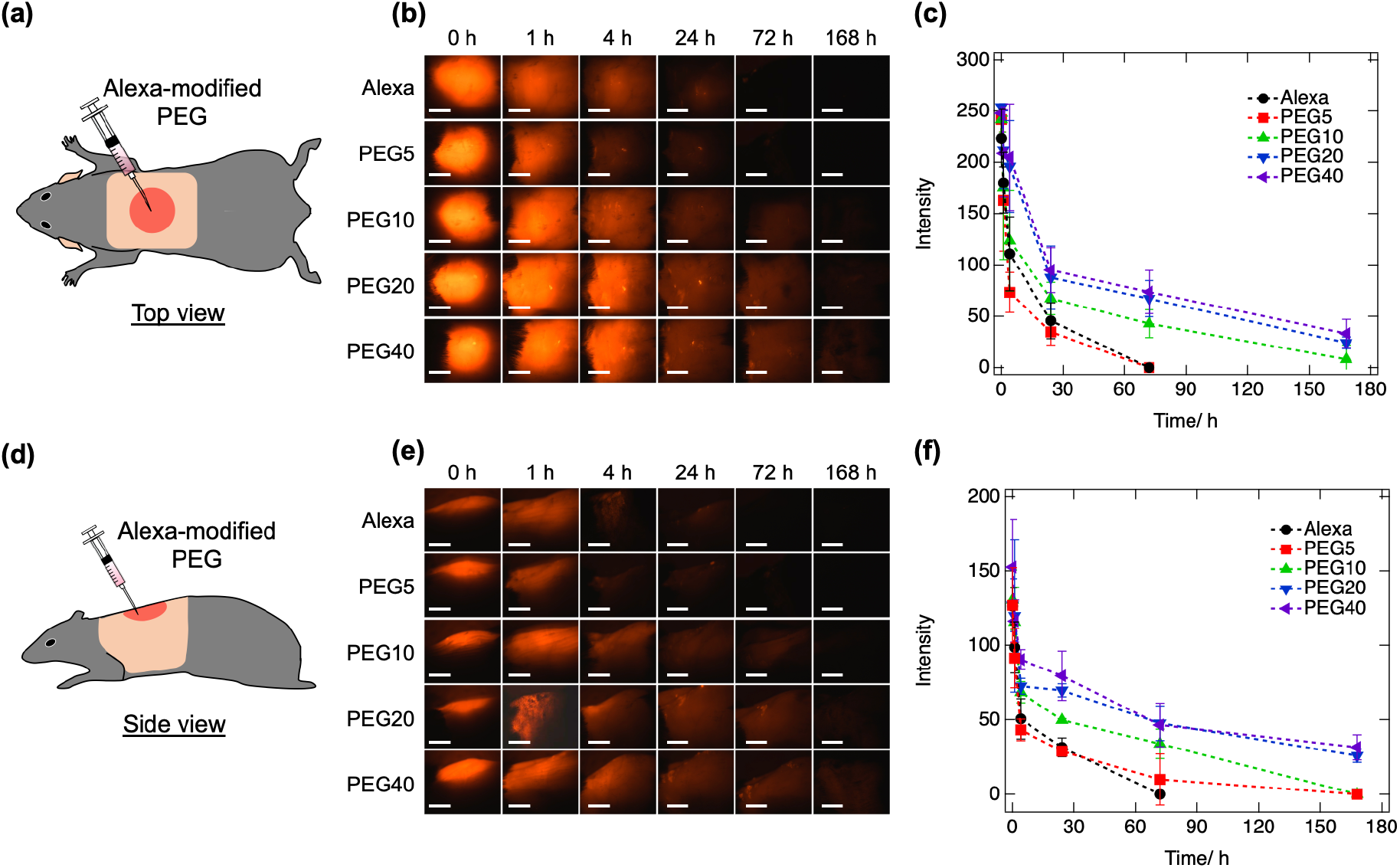
Diffusion behavior of Alexa-modified PEGs subcutaneously injected into the back of mice. **(a, d)** Diagram of injection site from the top (**a**) and side (**d**) of mice. **(b, e)** Fluorescence images from the top (**b**) and side (**e**) of mice at 0, 1, 4, 24, 72, and 168 h post-injection with Alexa-modified PEGs solutions. Scale bar: 5 mm. **(c, f)** Changes in fluorescence intensity over time from the top (**c**) and side (**f**) of mice after injection of Alexa (black circle), Alexa-modified PEG5 (red square), PEG10 (green triangle), PEG20 (blue inverted triangle), and PEG40 (purple left-arrowed triangle). Dashed lines between plots are shown as a guide.

Deep adipose tissue beneath the injection site (extending back to the axillar) was dissected as subcutaneous tissues toward regional lymph nodes (**Figure 3a**). Fluorescence intensity changes over time were evaluated with samples obtained at 0, 1, 4, 24, 72, and 168 h after injection (n = 3 each; **Figure 3b** and **c**). Alexa-labeled PEG was detected within deep adipose tissues collected just after injection (0 h), indicating that subcutaneously injected PEG was distributed into the deep tissues away from the injection site. Time-wise analyses illustrated differential findings between PEG with different *M*_w_. Fluorescent signals with PEG5 disappeared promptly, similar to the Alexa-only solution. Signals with PEG10 extended evenly to the whole tissues and consistently decreased over the 168 h of observation, while the reduction of the spread of fluorescence signals of PEG20 and PEG40 stagnated within 168 h of observation. Thus, diffusion of PEG from adipose tissue could be differentially classified into the following three categories: (i) fast solution-like diffusion (PEG5), (ii) gradual diffusion (PEG10), and (iii) resistant diffusion (PEG20 and PEG40).

**Figure 3.**
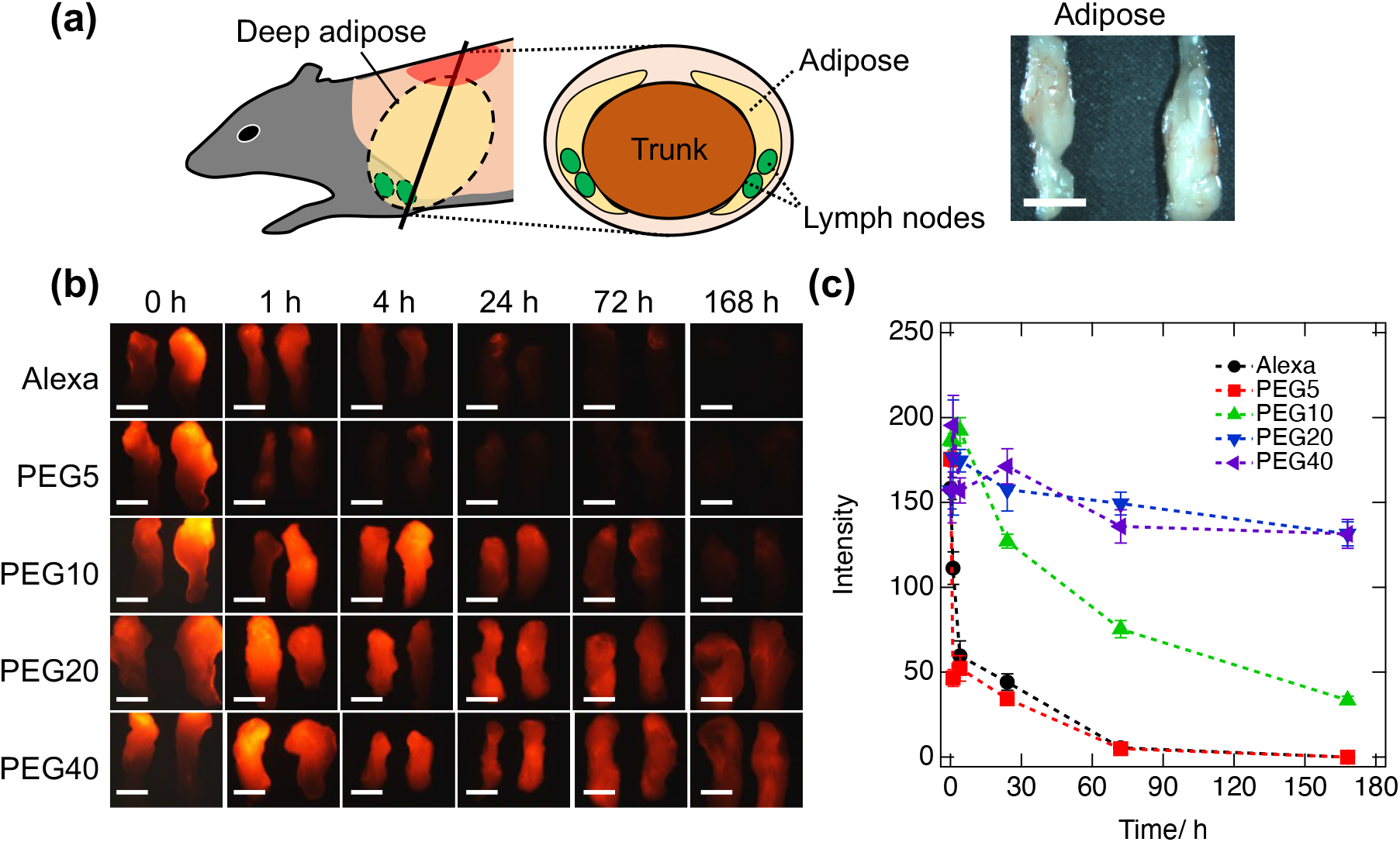
Distribution of Alexa-modified PEGs in the deep adipose tissues of mice. **(a)** Diagram of dissected subcutaneous deep adipose tissues toward regional lymph nodes. Photographs show representative deep adipose tissues. Scale bar: 5 mm. **(b)** Fluorescence images of deep adipose tissues at 0, 1, 4, 24, 72, and 168 h post-injection with Alexa and Alexa-modified PEGs. Scale bar: 5 mm. **(c)** Time-wise fluorescence intensity changes in the adipose tissues after injecting Alexa (black circle), Alexa-modified PEG5 (red square), PEG10 (green triangle), PEG20 (blue inverted triangle), and PEG40 (purple left-arrowed triangle). Dashed lines between plots are shown as a guide.

Detailed PEG diffusion in subcutaneous tissues was further analyzed in a lump of adipose tissues and axillary lymph nodes collected 4 h after PEG10 injection (gradual diffusion pattern; **Figure 4a**). With dispersed distribution throughout the tissues, localized PEGs were confirmed in lymph nodes (**Figure 4b**). A portion of subcutaneously injected PEGs migrates into regional lymph nodes. Similar to reports for macromolecules,^26,28^ the lymphatic vascular plays a significant role in the migration of subcutaneously injected PEGs into the blood circulatory system.

**Figure 4.**
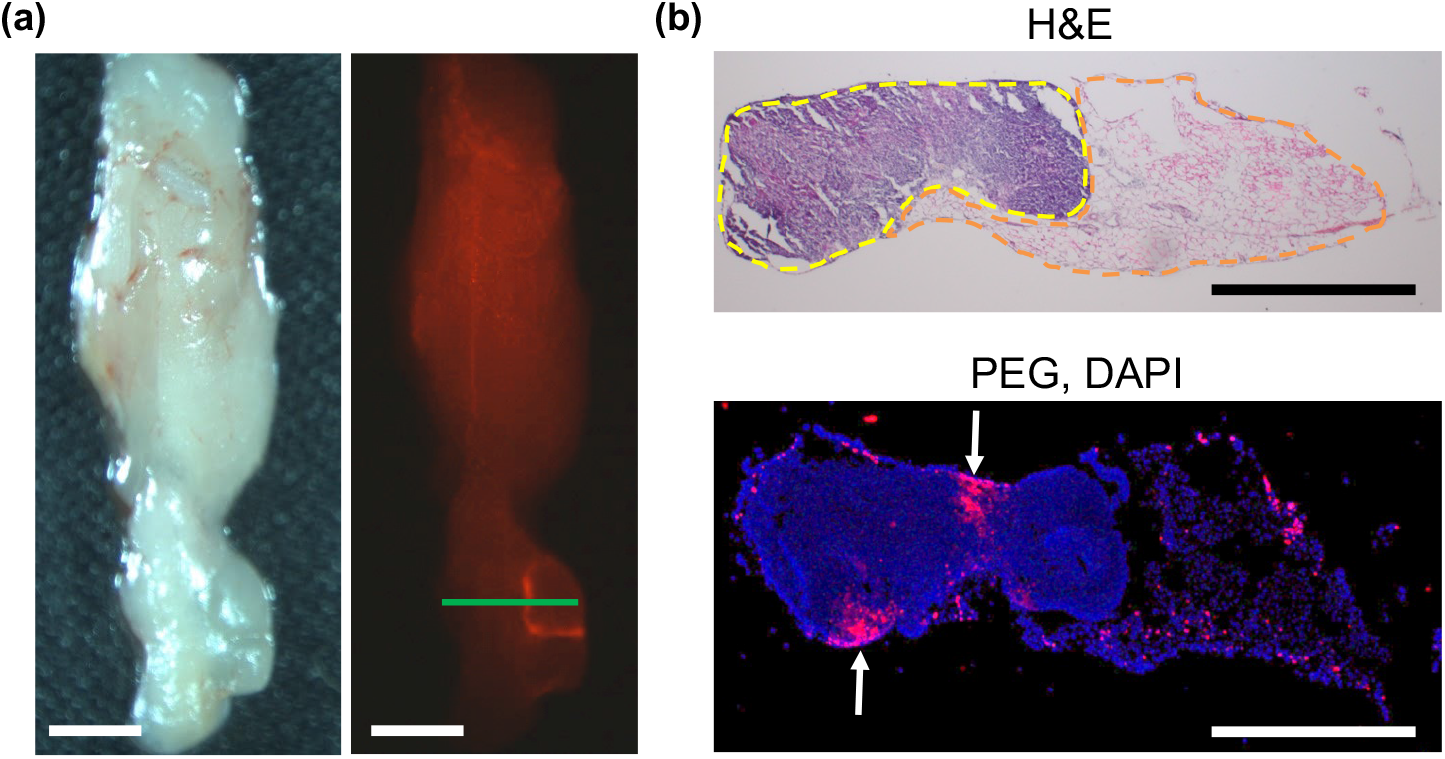
Representative images of adipose 4 h after PEG10 injection. **(a)** Representative bright vision (left) and fluorescence image (right) of deep adipose tissues. A solid green line indicates the position of the section. Scale bar: 2 mm. **(b)** Slice image of deep adipose samples stained with hematoxylin and eosin (H&E; top) and visualized by PEG and DAPI (bottom). In H&E staining, yellow and orange solid lines show the lymph nodes and deep adipose tissue, respectively. In the fluorescence image, red and blue show PEG-modified with Alexa and DAPI-stained nuclei, respectively. White arrows show PEGs accumulated in lymph nodes. Scale bar: 1 mm.

It is well-known that circulating macromolecules such as PEG show differential organ distribution depending on their *M*_w_.^8,12,13^ Especially in the kidney, macromolecules smaller than glomerular diameter migrate into the glomerulus, accumulate, and are excreted into the urine, while larger ones are hardly secreted.^29^ In the current study, the hydrodynamic diameter of PEG10 corresponded to the size of the known glomerular filtration limit (3–5 nm).

To observe the systemic distribution profiles of PEG after subcutaneous injection, over time fluorescence intensity changes in the heart, lung, liver, and kidney were evaluated at 0, 1, 4, 24, 72, and 168 h post-injection (n = 3 each; **Figure 5a,b**, and **Figure S1**). Consistent with a previous report, there was an apparent difference between the kidney and other organs.^8^ In the kidney, the signal intensity of PEG5 and PEG10 (peaked at 1–4 h) was higher than those of PEG20 and PEG40, while signals of PEG20 and PEG40 (peaked at 24 h) were higher in the heart, lung, and kidneys (**Figure 5b**).

**Figure 5.**
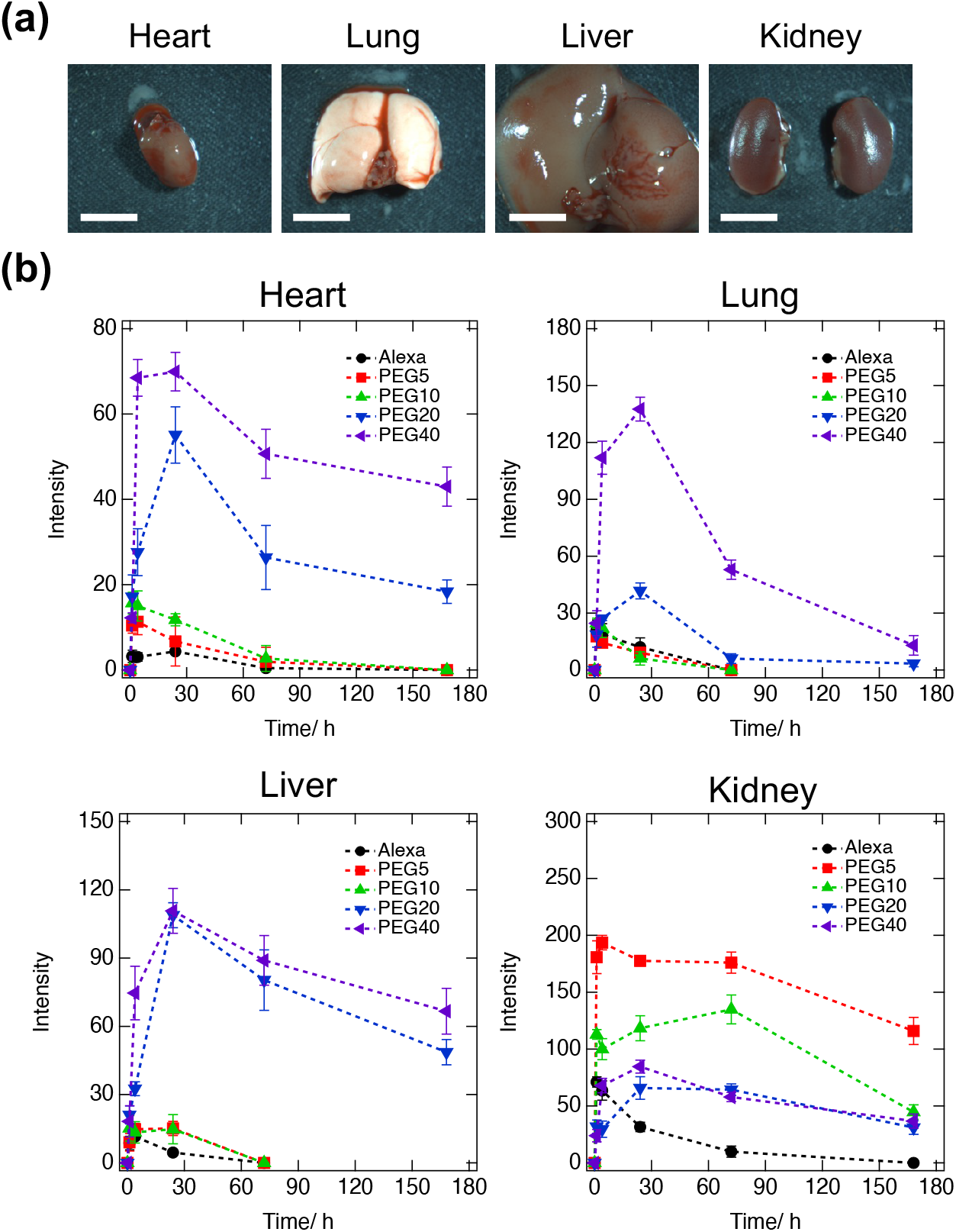
Biodistribution of Alexa-modified PEGs injected subcutaneously into the back of mice. **(a)** Representative photos of the heart, lung, liver, and kidney. Scale bar: 5 mm. **(b)** Fluorescence intensity changes over time of Alexa (black circle), Alexa-modified PEG5 (red square), PEG10 (green triangle), PEG20 (blue inverted triangle), and PEG40 (purple left-arrowed triangle) distributed in the heart, lung, liver, and kidney. Dashed lines between plots are shown as a guide.

To evaluate the biodistribution and clearance behavior of PEGs in the kidney, urine collected from the bladder underwent fluorescence analysis (**Figure 6a**). Although a reliable quantitative evaluation was difficult, fluorescence signals could be detected in Alexa, PEG5, and PEG10 samples at earlier time points (1–4 h). Further histological analyses of the kidney samples obtained 4 h after PEG10 injection revealed highly accumulated fluorescence in glomeruli (**Figure 6b**).

**Figure 6.**
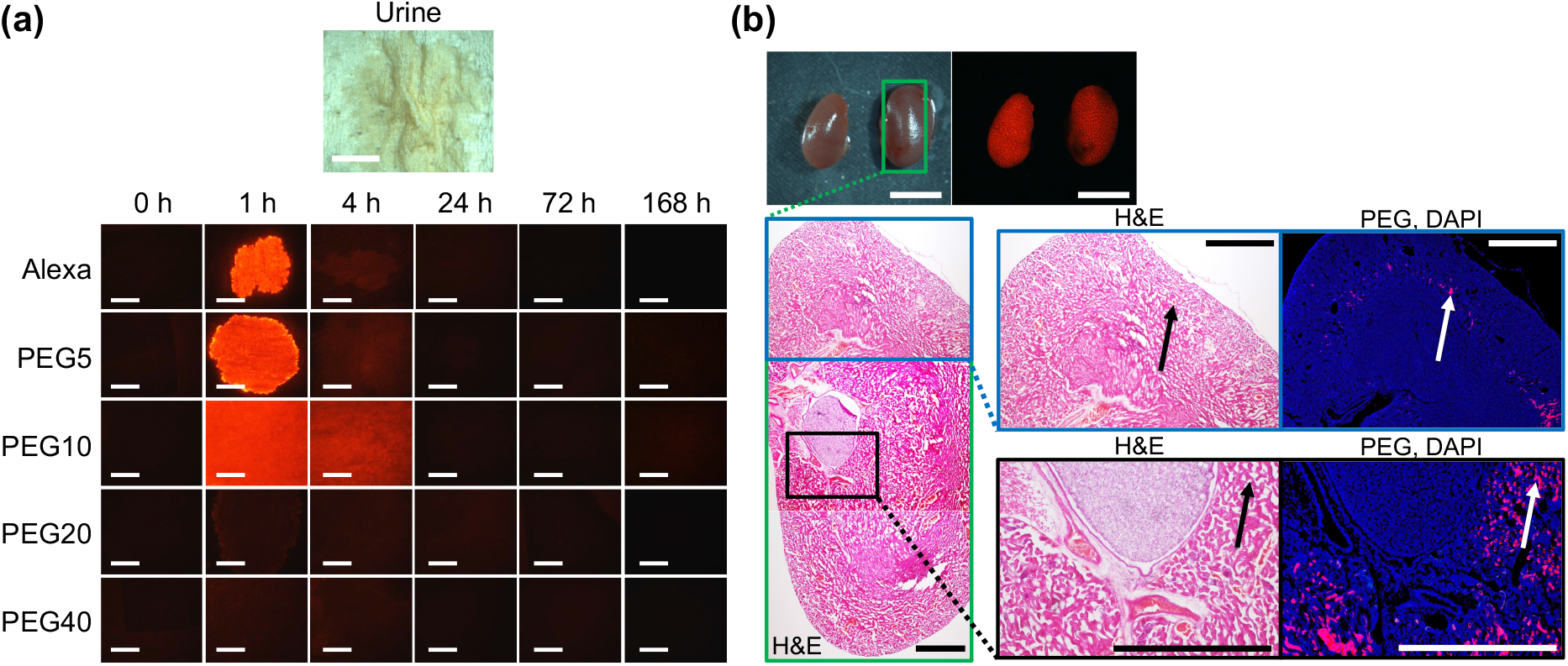
Clearance behavior of PEGs in the kidney. **(a)** Photos and fluorescence images of urine collected from the bladder at 0, 1, 4, 24, 72, and 168 h after injection with Alexa and Alexa-modified PEGs solution. Scale bar: 5 mm. **(b)** Histological images of the kidney 4 h after injection with PEG10. The location of H&E staining and sliced fluorescence images are indicated by green, blue, and black lines. Scale bar: 5 mm in the photographs and fluorescence images and 1 mm in histological images. The renal cortex glomerulus is indicated using black and white arrows.

In the current study, time-wise distribution of subcutaneously injected PEGs in the skin, subcutaneous deep adipose tissue, lymph nodes, and distant organs such as the heart, lung, liver, and kidney were comprehensively characterized. Our findings are consistent with the hypothesis that a portion of subcutaneously injected PEGs is collected into the circulation system via lymphatic nodes and distributed to distant organs. PEG distribution in distant organs is considered to be in equilibrium with the circulatory system, and PEGs in the circulatory system are defined by influx from subcutaneous tissues and excretion from the kidney and liver. Based on our analyses, PEGs with lower *M*_w_, such as PEG5 and PEG10, disappear from the skin and subcutaneous tissues and are delivered to and possibly excreted dominantly in the kidney. The entire process was rapid. On the other hand, PEGs with higher *M*_w_, such as PEG 20 and PEG40, resided for longer periods in the skin, subcutaneous tissues, and distant organs other than the kidney.

Based on these experimental results, the following two essential points were made. First, the *M_w_* of PEG should be carefully selected according to the biomedical application. PEG with *M*_w_ ≥ 40 kg/mol accumulates in the liver and is consequently advantageous for drug delivery carrier applications, which require long-term localized action, such as Pegasys (also called Peginterferon α-2a).^30^ On the other hand, ethylene glycol units are cleaved by *in vivo* oxidative biodegradation on a long time scale, resulting in the generation of toxic radicals.^31^ Thus, although PEG has unique biocompatibility, it is advisable to utilize PEGs with suitable *M*_w_. In particular, for the development and use as hydrogel scaffolds that require complete disappearance after functionalization as biomaterials, it may be reasonable to select PEGs with a *M*_w_ below 10 kg/mol as precursors of hydrogels because such PEGs show gradual diffusion and clearance, primarily in the kidney.

Second, fluorescent labeling and detection are more advantageous than previously described methods. The present detection method is superior to existing ones for real-time imaging. To observe the clearance behavior of intravenously injected polymers, radiolabeled isotopes such as ^14^C and ^125^I have been introduced into PEG molecules instead of fluorescence-labeled PEG molecules.^8,13^ However, the real-time clearance behavior could not be observed owing to technical difficulties, thereby limiting the real-time evaluation of subcutaneous distribution. Furthermore, introducing such isotopes is not ideal because this requires a multi-step synthetic process,^13^ and isotopes show toxicity.^32,33^ The fluorescence labeling in this study is reasonable in terms of the simple preparation method and low toxicity of the fluorescence dye. On the other hand, multiple fluorescence imaging is also one of the important properties of our study, allowing us to clearly detect PEG localization in distant organs by detecting various fluorescence, such as cells and polymers (as shown in Figure 4b and 6b).

Nevertheless, despite these advantages, this red fluorescence imaging is unsuitable for real-time observation in deep tissues because it is difficult to detect fluorescence due to high tissue absorption and autofluorescence.^34^ For this reason, the *M*_w_-dependent clearance pathway was not fully investigated, as the fluorescence intensity could not be observed in the excrement. Such a clearance pathway is essential for the biocompatible evaluation of polymers, where scaffolds are implanted to regenerate defective tissues. In this case, employing near-infrared (NIR) fluorescence may allow real-time detection of PEGs because of relatively minimum tissue absorption and autofluorescence. Therefore, although special equipment is required, multiple methods, such as isotope and NIR observation, should be applied in the future to comprehensively detect PEG localization.

In summary, Alexa-modified PEGs subcutaneously injected into the backs of mice showed *M*_w_-dependent diffusion in the skin, biodistribution to distant organs, and clearance behavior. PEGs with *M*_w_ ≤ 10 kg/mol gradually diffused in the subcutaneous tissue, migrated to deep adipose tissue, and distributed to distant organs, mainly the kidney. On the other hand, PEGs with a *M*_w_ ≥ 20 kg/mol stagnated in the skin and were mainly delivered to the heart, lungs, and liver. The findings of this study clarify the basis of the *in vivo* diffusion, biodistribution, and clearance behavior of four-armed PEGs, which are currently extensively used as basic materials for producing clinically relevant biomaterials. Taken together, the observed diffusion, biodistribution, and clearance behavior clarified the fate of four-armed PEGs, thus providing an essential reference for synthesizing and implementing PEG-based biomaterials.

## Associated Content

Additional experimental details, materials, and methods; results of tests conducted to evaluate fluorescence intensity; photos of collected tissues; this material is available free of charge via the Internet at http://pubs.acs.org.

## Corresponding Authors

* Shohei Ishikawa

* Takamasa Sakai

* Masakazu Kurita

## Author Contributions

S.I. and M.K. designed the study. S.I., M.K., J.S., L.C.Y., and K.K. performed all the experiments. S.I., M.K., T.K., and M.N. performed data acquisition and/or analysis. All authors drafted and reviewed the manuscript. Administrative, technical, and supervisory tasks were handled by S.I., M.K., and T.S.

## Funding Sources

This work was supported by the Japan Society for the Promotion of Science (JSPS) Fellows (Grant No. 20J01344), a Grant-in-Aid for Young Scientists (Grant No. 21K18063) to S.I., JSPS Fellows (Grant No. 21J10828), Transformative Research Areas grant number 20H05733 to T.S. This study was also supported by the Japan Science and Technology Agency (JST) CREST Grant number JPMJCR1992 to T.S., and JST Moon-shot R&D grant number 1125941 to T.S. Grant-in-Aid for Challenging Research (Pioneering; Grant No. 20K20609) to M.K., and AMED Moon-shot R&D (Grant No. 21zf0127002h0001) to T.S. and M.K.

## Conflict of Interest

The authors declare no conflict of interest.

## Table of Contents

Four-armed poly(ethylene glycol) (PEG) is an essential polymer utilized to prepare hydrogels called tetra-PEG gels, which are very important biomaterials in the field of tissue engineering. In this study, we observe the basic clearance behavior of four-armed PEGs subcutaneously injected into the back of mice by means of red fluorescence (Alexa)-labeled PEG. Our findings are expected to serve as an essential indicator for synthesizing biomaterials *in vivo*.

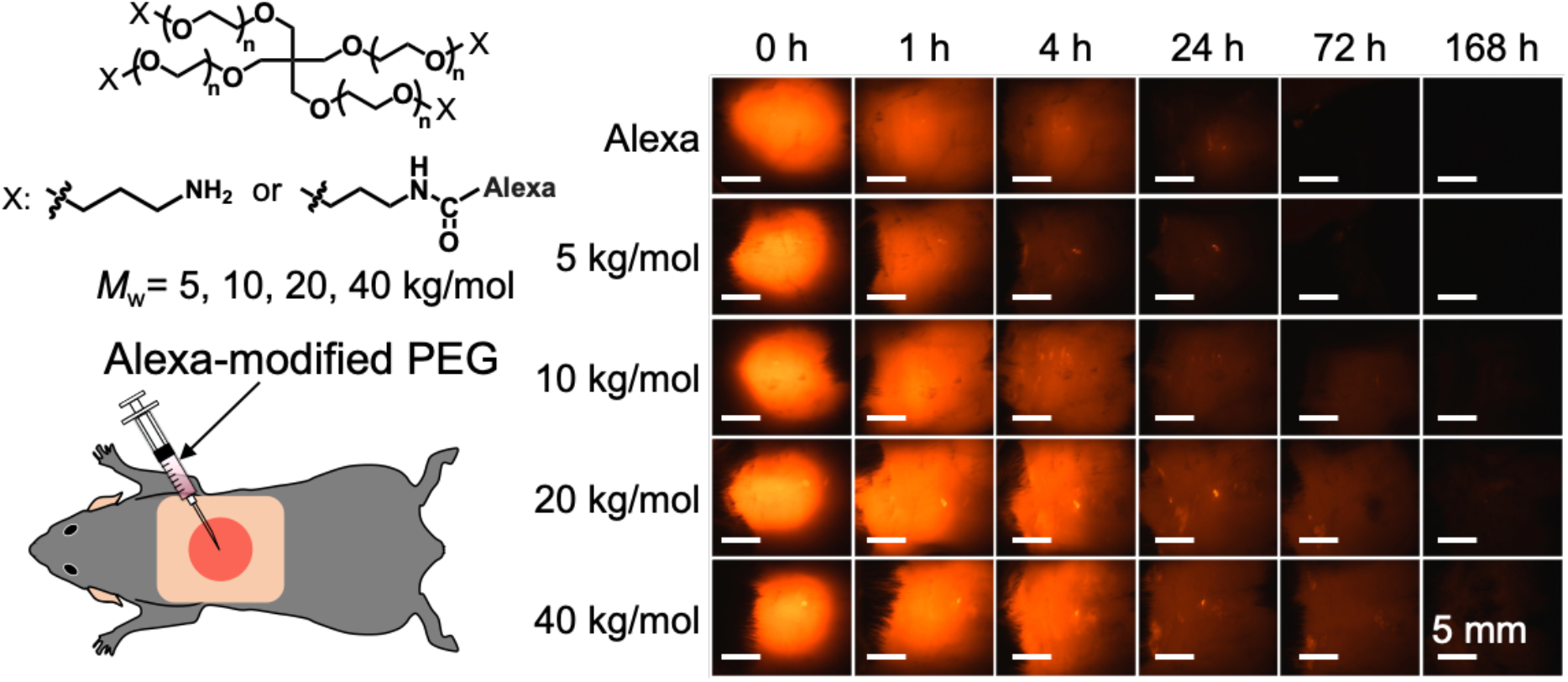

## Materials and Methods

### Materials

Propylamine-terminated tetra-armed PEG (Tetra-PEG-PA) with molecular weight (*M*_w_) = 5, 10, 20, and 40 kg/mol, also marketed as PTE-50-PA, PTE-100-PA, PTE-200-PA, and PTE-400-PA, were purchased from NOF CORPORATION (Tokyo, Japan), and used without further purification. Dulbecco’s phosphate-buffered saline (D-PBS) and Alexa Fluor™ 594 NHS Ester (Succinimidyl Ester) (Alexa-NHS) was purchased from Thermo Fisher Scientific (Massachusetts, USA). 4% paraformaldehyde phosphate buffer solution (PFA) was purchased from FUJIFILM Wako Pure Chemical Corporation (Tokyo, Japan). Alexa-NHS was dissolved with DMSO to get the concentration of 1 mg/mL. 12-well culture plate, 12-well cell culture insert (pore size of 1.0 μm), and 96-well plate were purchased from (Becton, Dickinson and Company, NJ, USA). MCR 301 rheometer (Anton Paar, New South Wales, Austria), and ARVO™ X3 microplate reader (PerkinElmer, Inc., Massachusetts, USA), Milli-Q was used as water in this study.

### Preparation of Alexa-modified PEGs

Tetra-PEG-PA was dissolved in D-PBS to obtain the concentration of 100 g/L. Alexa-NHS was added to prepared PEG solution to 0.1% vs. primary amine, and the mixture was incubated at 25 °C for 24 h under dark condition. The incubated PEG solution was then dialyzed against water for 24 h to remove unreacted Alexa-NHS molecules, and freeze-dried to obtain Tetra-PEG-PA functionalized partly with Alexa that were referred to as Alexa-modified PEG.

### Viscosity measurement of Alexa-modified PEGs

Tetra-PEG-PA and Alexa-modified PEG were separately dissolved in D-PBS to obtain the concentration of 10 g/L. The viscosity (*η*_PEG_) in steady state was measured at 25 °C as a function of the shear rate (*γ*) ranging from 0.027 to 2.7 s^-1^. The measurements were performed with the cone-plate fixture (diameter: 50 mm, cone angle: 4°). The viscosity (*η*) relative to that of PBS (*η_s_*) was calculated as *η* = *η*_PEG_/*η*_s_.

### In vitro release of Alexa-modified PEGs

The release model, which is set on 12-well culture plate with 12-well cell culture insert with a pore size of 3.0 μm, was designed. First, 200 μL and 1200 μL of D-PBS were added to the insert and well plate, respectively. Then, 100 μL of Alexa-modified PEG solution, which is prepared with D-PBS to obtain the concentration of 10 g/L, was added to the insert. At 25 *°C* under dark condition, 1200 μL of sampling solution in the well plate was collected, and 1200 μL of D-PBS was added at every time point. After that, 300 μL of sampling solution was added to 96-well plate. The concentration of sampling solution was evaluated by measuring fluorescence intensity (*λ_ex_* = 580 nm, *λ*_em_ = 590 nm) using a microplate reader. The release rate was evaluated as a percentage compared with total amount of Alexa-modified PEG.

### Injection of Alexa-modified PEGs into the back of mice

C57BL/6 mice, 4-week-old females, were purchased from Nippon Bio-Supplement Center (Tokyo, Japan) and used for animal experiments. Under inhalation anesthesia with Isoflurane, the upper body was shaved circumferentially with a razor blade. Then, 100 μL of Alexa-modified PEG solutions were injected subcutaneously around the center of both shoulder blades in the middle of the back.

After each time period, fluorescence intensity was evaluated. Each mouse was given inhalation anesthesia, and euthanized by cervical dislocation. To evaluate the diffusion behavior from the injection site, fluorescence intensity was observed from the top and side of the injection site by Axio Zoom (ZEISS, Germany). The heart, lungs, liver, kidneys, and subcutaneous deep adipose tissue contiguous to the bilateral axillae with the injection site as the upper edge were then carefully harvested. Furthermore, paper wetted by mouse incontinence was used as urine sample. Immediately after sampling of these organs, fluorescence intensity was evaluated by Axio Zoom. In each experiment, three animals were employed per condition.

### Histological staining

Dissected deep adipose tissue and kidney obtained 4 hours after PEG with *M_n_* of 10 kg/mol injection were fixed in 4% PFA for overnight and then incubated in 30% sucrose. Frozen samples were then prepared with optimal cutting temperature compound (O.C.T. Compound®, Sakura Finetek, Japan), and cut into 12 μm slices using a cryostat. These slices were then fixed in 4’,6-diamidino-2-phenylindole (DAPI) mounting media (Fluoromount-G^®^ with DAPI, Thermo Fisher Scientific, USA), and washed twice with PBS. On the other hand, these slices were also stained with hematoxylin and eosin (HE). These samples were observed by inverted microscope (Olympus, Japan).

## Supplementary Tables and Figures

**Table S1.**
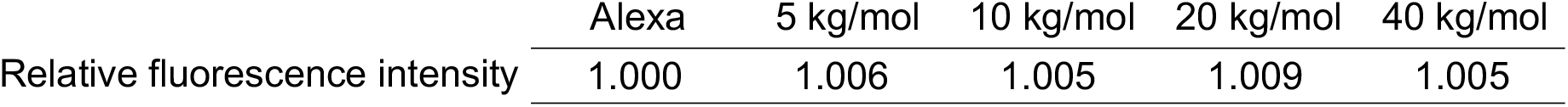
Relative fluorescence intensity of Alexa-modified PEGs.

**Figure S1.**
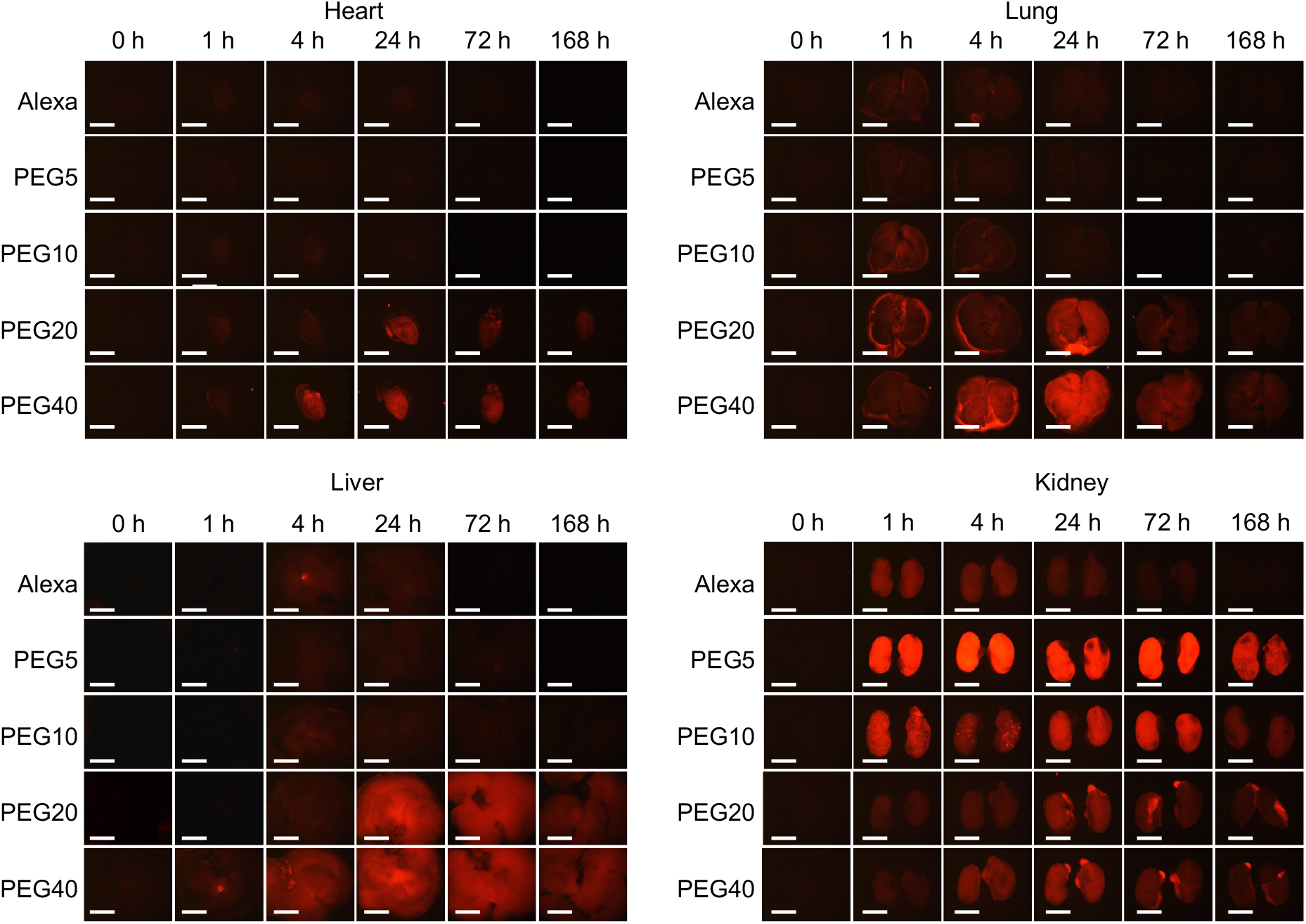
Fluorescence images of heart, lung, liver, kidney, adipose, and urine. Scale bars show 5 mm.

## Notes

### Competing Interest Statement

The authors have declared no competing interest.

